# smBEVO: A computer vision approach to rapid baseline correction of single-molecule time series

**DOI:** 10.1101/2021.11.12.468397

**Authors:** Khue Tran, Argha Bandyopadhyay, Marcel P. Goldschen-Ohm

## Abstract

Single-molecule time series inform on the dynamics of molecular mechanisms that are occluded in ensemble-averaged measures. Amplitude-based methods and hidden Markov models (HMMs) frequently used for interpreting these time series require removal of low frequency drift that can be difficult to completely avoid in real world experiments. Current approaches for drift correction primarily involve either tedious manual assignment of the baseline or unsupervised frameworks such as infinite HMMs coupled with baseline nodes that are computationally expensive and unreliable. Here, we develop an image-based method for baseline correction using techniques from computer vision such as lane detection and active contours. The approach is remarkably accurate and efficient, allowing for rapid analysis of single-molecule time series contaminated with nearly any type of slow baseline drift.

## Introduction

Single-molecule time series inform on the dynamic conformational changes that underlie molecular mechanisms. Their analysis involves assignment of trajectories amongst discrete states that reflect changes in amplitude or duration of a piecewise continuous signal. These trajectories are otherwise occluded in ensemble measures due to averaging over asynchronous behavior in many molecules. Such data is provided by a variety of increasingly common experimental modalities including single-channel electrophysiology, fluorescence imaging, magnetic tweezers, and atomic force microscopy (Goldschen-Ohm et al., 2016; Mortensen & Smart, 2007; Juette et al., 2016; Ma et al., 2017; McKinney et al., 2006; Ulbrich & Isacoff, 2007; White et al., 2021; Gu et al., 2017). Ideally, experiments are designed to minimize baseline drift of the recorded signal independent of signal changes arising from the behavior of the system of interest. However, signal drift can arise from a multitude of complications such as focal drift of the imaging objective, electrical noise, mechanical stress and vibrations, for example during perfusion and solution exchange, changes in temperature, bleaching of background fluorescence, movement of molecules across polarized or nonuniform excitation fields such as for total internal reflection fluorescence (TIRF) microscopy, fluorophore photo dynamics, instability of laser excitation sources, and fluidity of the sample where applicable (e.g. lipid membranes), and thus can be difficult to eliminate completely in real world experiments (Carter et al., 2007; Colomb et al., 2017; Holden et al., 2010; Lelek et al., 2021; Nugent-Glandorf & Perkins, 2004; Smith et al., 2003; Vivaudou et al., 1986; Woodside et al., 2006).

Analysis of single-molecule time series relies on idealization of the noisy measured signal into discrete dwells in distinct states. Idealization methods such as change point detectors or HMMs rely heavily on the amplitude of the signal to identify periods in distinct states (Watkins & Yang, 2005; Shuang et al., 2014; White et al., 2020; Loeff et al., 2021; Riessner et al., 2002; Schultze & Draber, 1993; Bronson et al., 2009; Qin et al., 2000). These approaches are severely compromised by baseline drift because they assume changes in signal amplitude reflect changes in state rather than a change in baseline. Thus, accurate analysis of single-molecule time series requires that baseline drift be removed.

A common supervised approach to correct for drift involves manual placement of baseline nodes and subsequent subtraction of a time varying baseline interpolated along the nodes (Bruno et al., 2013; Nicolai & Sachs, 2013; Talukder & Wollmuth, 2011; Wang et al., 2016). This approach can provide excellent results, but is highly time consuming, eventually becoming intractable for very large datasets that are increasingly common for high-throughput experimental paradigms. Node placement can be automated by maximizing the likelihood of an HMM and a drifting baseline interpolated along a set of variable node positions (Syed et al., 2010; Venkataramanan & Sigworth, 2002; Y. Zhang et al., 2016). However, global analysis with a specific model is often not feasible for datasets with per-molecule variation in behavior or state emission and noise amplitudes. Variation in behavior is common in large high-throughput datasets where typically only a subset of molecules exhibits the behavior of interest. Variation in signal-to-noise is typically observed in camera-based imaging approaches due to spatial nonuniformities in the optical pathways and/or illumination (Brunstein et al., 2014). Dynamic variation in signal properties due to fluorophore photo dynamics or motion within polarized or steeply varying excitation fields such as for TIRF microscopy can also challenge analysis with specific HMMs (Dempsey et al., 2009; Levene et al., 2003; Oheim et al., 2019; Stennett et al., 2015; White et al., 2020). Finally, the requirement for postulation of a specific mechanistic model can be a limitation when the underlying mechanism is either unknown or not well constrained. Thus, model-free approaches are desirable. Local jump detection has been used to estimate drift but requires the drift to be essentially nonexistent between any two consecutive jumps which will not be the case in general (Raillon et al., 2012; Vivaudou et al., 1986). Other model-free approaches are often limited by requirements such as uniform step sizes that are appropriate in only specific circumstances (Bruno et al., 2013). For camera-based imaging, neighboring pixel intensities can potentially be used to approximate a time varying baseline so long as it is due to variation in the optical pathway as opposed to local variation of at the molecule (Möckl et al., 2020; Preus et al., 2015). Deep learning approaches can potentially handle slow baseline drift but require pre-idealized training data (Celik et al., 2020). In particular, two methods based on infinite HMMs (iHMMs) or minimum description length (MDL) allow for unsupervised model-free analysis (Ouqamra & Bouilly, 2019; Sgouralis et al., 2018). However, as discussed below these two methods are unreliable for typical single-molecule time series data.

Thus, a model-free approach that rapidly and reliably corrects for baseline drift in general single-molecule time series with potentially nonuniformly spaced levels is needed. Here, we implement frameworks from image analysis including lane detection algorithms for automated driving and active contours for boundary detection to develop a tool that quickly and accurately removes baseline drift from single-molecule time series with minimal user dependence.

## Results

### smBEVO: single-molecule baseline estimation via visual optimization

We leverage methods from computer vision to develop an image-based approach for estimation of time varying baseline drift in single-molecule time series. The approach, smBEVO, depends primarily on only a few parameters that can be intuitively estimated and are likely to be shared amongst all series under similar experimental conditions. Throughout, we use the variables *x* and *y* to denote the series propagation (e.g. time) and signal amplitude, respectively (Fig. 1a). Although we establish the approach in the context of single-molecule time series, it is viable for any piecewise continuous signal with a discrete set of reasonably separated amplitude levels.

**Fig 1.**
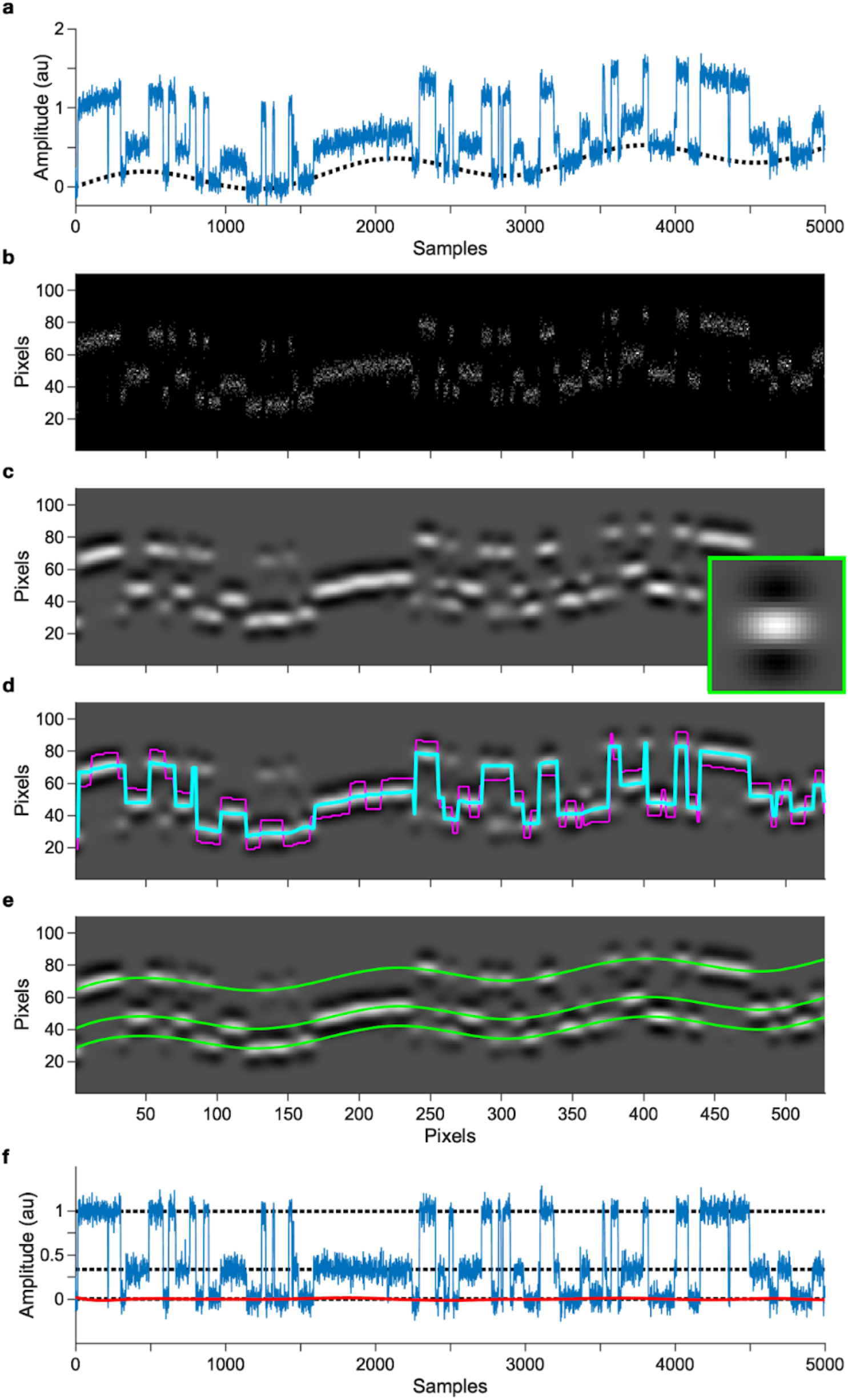
smBEVO baseline correction workflow. **a**, Simulated time series for a cyclic 3-state model with three distinct intensity emissions (one per state) at amplitudes of 0, 1/3 and 1 (arbitrary units) with added linear and sinusoidal baseline drift (dotted). **b**, Image representation of the series data using *σ*_*x*_= 38 samples and *σ*_*y*_= 0.11 au with default *s*_*y*_= *s*_*y*_= 4 pixels. **c**, Filtered image after convolving the raw data image with a selective oriented filter kernel (inset). **d**, Initial idealized trajectories based on the sequences of maximum (cyan) or minimum (magenta) intensity pixels in each column and a smoothing factor of *λ*=10. **e**, Parallel levels (green) derived from segmentation of the initial idealized trajectories above and a minimum level separation *δ*_*y*_= 0.3 au. **f**, Baseline-corrected series data after subtracting the lowest identified level following its interpolation from image to data coordinates. Dotted lines indicate identified levels. Red line is the residuals between simulated and estimated baselines.

The series data is first represented as a grayscale image at a resolution determined by user defined scale factors *σ*_*x*_ and *σ*_*y*_ and a specified number of pixels per scale factor set to *s*_*x*_= *s*_*y*_= 4 pixels (see Methods). The image is obtained as a normalized two-dimensional histogram of the series (*x, y*) data points with bin widths in each dimension of approximately *σ*_*x*_/*s*_*x*_ and *σ*_*y*_/*s*_*y*_ (Fig. 1b, see Methods). Finally, the image is filtered by convolving with a two-dimensional selective oriented Gaussian kernel (Fig. 1c; Eqs. 1-2). This filter is tuned to detect horizontal bands of high intensity and was previously used as part of a lane marker detection algorithm for computer vision-based approaches to automated driving (Aly, 2008). For single-molecule time series, this filter effectively blurs out rapid signal fluctuations due to noise and accentuates the y-axis separation between piecewise continuous segments of differing amplitudes while also blurring together nearby segments of similar amplitude. The resulting filtered image provides an estimate of the primary amplitude levels in the data series and their variation in x due to slower changes in baseline. The filtered image and final baseline estimation depend heavily on *σ*_*x*_ and *σ*_*y*_, which can be reasonably estimated based on the data series. In practice, *σ*_*x*_ should be as large as possible to blur together periods where the baseline is undergoing minimal change, but not so large that significant changes in baseline are blurred together. Similarly, *σ*_*y*_ should be sufficiently large to smooth out fluctuations caused by noise, but small enough to not blur together amplitude levels that should be distinguished. Estimating *σ*_*y*_ as approximately one third of the minimum level separation to be resolved typically works well. We note that accurate identification of a slowly drifting baseline does not necessarily require resolution of every level or event in the data, and in some cases blurring together closely spaced levels can still provide good baseline estimation.

After filtering, an initial denoised trajectory in image coordinates is estimated as the sequence of pixels with maximum intensity in each column (one pixel per column) (Fig. 1d). This denoised trajectory has the very useful property of providing a highly filtered version of the data series while largely maintaining sharp jumps between amplitude levels that are otherwise blurred out with conventional one-dimensional filters. The trajectory is divided into disjoint segments at changepoints identified by jumps exceeding a specified threshold of *s*_*y*_/2 in either the maximum intensity trajectory or its conjugate minimum intensity trajectory (see Methods). Optionally, each segment is smoothed using a piecewise cubic spline with spline segments of length approximately *λ*/*s*_*x*_ (may be slightly shorter to fit an integer number of spline segments), where *λ* is a specified smoothing parameter (larger values for more smoothing, zero implies no smoothing).

A set of distinct amplitude levels is identified by first assigning an initial level to the longest segment. This level is then iteratively extended into the preceding and following neighboring segments. Each neighboring segment is assigned to an existing level if it is within a specified threshold of *δ*_*y*_ pixels at the join between segments. Otherwise, it is assigned to a new level. In the latter case, the new level is extrapolated back along all existing levels to maintain a constant separation with the nearest level. The existing levels are then extrapolated into the neighboring segment to maintain all level separations at the segment join. This is repeated until levels have been assigned across the entire image. Typically, a default value of *δ*_*y*_= 1.75*s*_*y*_ pixels works well, but increasing *δ*_*y*_ can mitigate the occurrence of spurious closely spaced levels due to noisy data. Optionally, the levels may be smoothed after each iterative addition of a segment with a piecewise cubic spline based on the smoothing parameter *λ* as described above. In addition to *σ*_*x*_ and *σ*_*y*_, the amount of smoothing as specified by *λ* can impact the resulting baseline estimation. Larger values of *λ* can be useful to mitigate spurious levels or drift arising from brief artifacts or larger noise fluctuations. Although levels as defined above are parallel within each segment, they may undergo nonidentical steps between segments. Thus, the levels are parallelized by replacing each level with a shifted version of the mean across levels, where the shift is determined so as to minimize the sum of squared errors with the original level. Supplementary Fig. 1 illustrates several steps during level construction.

Although appropriate choice of *σ*_*x*_, *σ*_*y*_ and *λ* are often sufficient for accurate baseline estimation, the set of identified levels can be further refined based on the image intensities by describing them as a set of parallel open-ended active contours, also called snakes. Snakes are widely used in image analysis to identify edges or contours around objects. They are flexible splines that are iteratively positioned on an image based on an external force field *F*_*ext*_ (typically a function of the image itself) and an internal energy expression *E*_*int*_ that is composed of terms that constrain the elasticity and curvature of the spline (Kass et al., 1988; Papari & Petkov, 2011). Here we set *F*_*ext*_ to the gradient of the image along the y-axis normalized to intensities between - 1 and 1. During each iteration, we define the external force on each column of each snake as the average gradient across each snake in that column. In this way, all the snakes will move together within each column to maintain their initial parallel level separations. Note that the forcefield is only applied along the y-axis and not along the x-axis. This results in snakes that converge on levels that maximize the overall intensity across all levels while enforcing a degree of smoothness via their internal energy terms. The parameters *α* and *β* scale the internal energy terms that constrain the allowable elasticity and curvature of the levels, respectively. Larger values of *α* and *β* result in smoother, less wiggly snakes. The rate of change of the nodes per iteration is scaled by the parameter *γ*. In practice, *γ*= 1 works well and likely does not need to be adjusted. The search terminates either after a specified maximum number of iterations or when the change in node positions falls below a tolerance threshold. In practice, 10-20 iterations are typically sufficient for the snakes to converge.

The baseline in image coordinates is set to the lowest amplitude level. Also, an idealized trajectory amongst levels is defined by projecting each segment onto the nearest level. This trajectory is likely to be a highly filtered version of the true ideal trajectory in the raw data but may nonetheless still be useful for preliminary analysis and screening of molecules of interest. The baseline, levels, and idealized trajectory are converted back to the original data coordinates by interpolating between pixel centers. For simplicity and speed, we use linear interpolation. Finally, the estimated baseline is subtracted from the raw data series to obtain a series that is ready to be analyzed by any available amplitude-dependent method such as step detectors or HMMs (Fig. 1e).

In practice, selection of appropriate values for *σ*_*x*_ and *σ*_*y*_ are often all that is needed for good baseline estimation. If this does not yield a reasonable result, then adjusting the smoothing parameter *λ*, and in some cases the minimum level separation *δ*_*y*_, are usually sufficient. Finally, the identified levels can be further refined based on the image gradient with active contours by adjusting *α* and *β* for desired flexibility.

### Validation of smBEVO on simulated data

To validate smBEVO we simulated single-molecule time series for several Markov models composed of 2-5 states with distinct emission amplitudes and either bidirectional or unidirectional transitions (Fig. 2a, see Methods). Gaussian noise was added to generate noisy series at various signal-to-noise ratios (SNRs), where SNR is defined as the ratio of the average separation between neighboring amplitude levels and the standard deviation of the added noise. We note that for nonuniform level separations in the cyclic three-state model some events will have either higher or lower SNR than the specified average. Finally, one of five different types of baseline drift was added to each time series to simulate various empirically observed drifts: no drift, linear drift, sinusoidal drift, linear + sinusoidal drift, or exponential decay of the baseline (see online methods for details). For each unique combination of model, SNR, and drift type we simulated 100 time series each with a length of 5000 samples and applied smBEVO to estimate the simulated baseline drift.

**Fig 2.**
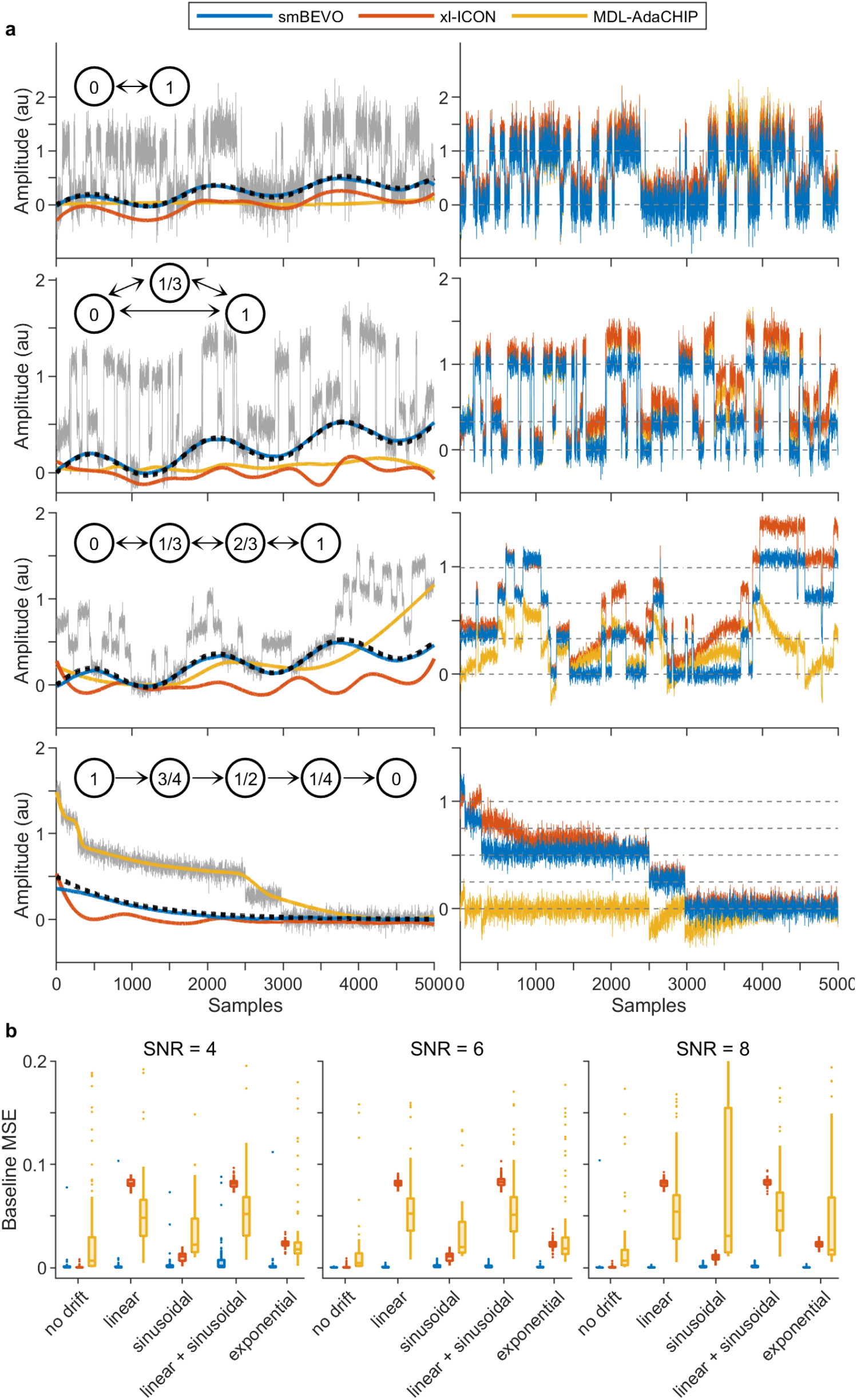
Comparison of smBEVO with xl-ICON and MDL-AdaCHIP. **a**, Simulated noisy time series for four different mechanisms (insets, see Methods) with added baseline drift (dotted) overlaid with baseline estimations for smBEVO, xl-ICON or MDL-AdaCHIP (left). Baseline subtracted time series for each method are shown to the right overlaid with the drift-free state emission levels for the associated mechanism (dashed). **b**, Summary box plots of the mean squared error (MSE) between simulated and estimated baselines for the four-state linear model with several types of applied baseline drift and signal-to-noise ratios (SNRs). See Supplementary Figs. 2-9 for additional examples and summaries for all tested models, drift types and SNRs. See Supplementary Tables 1-4 for smBEVO parameters in all conditions.

For each set of 100 series, we manually identified parameters for smBEVO that resulted in good baseline estimation of a handful of series, and thereafter applied these parameters uniformly to all 100 series (see online methods for details). This approach reflects the ideal workflow for a large dataset: first identify appropriate parameters under user supervision, and then use these parameters for unsupervised baseline correction of the entire dataset. In nearly all cases smBEVO provided an excellent estimation of the baseline even for fairly extreme baseline drift/wobble (Fig. 2a). Fig. 2b summarizes the mean squared error (MSE) between simulated and estimated baselines for all series simulated using the linear four-state model at several SNRs and all drift types. Example traces for each model at each SNR and with each drift type, as well as summary plots of MSE between simulated and estimated baselines for all tested time series are shown in Supplementary Figs. 2-9.

Overall baseline estimation was both visually and quantitatively compelling across models, SNRs and drift types. The only exception where smBEVO was less compelling is for the three-state cyclic model with sinusoidal drift at the lowest tested average SNR=4. We note that given the nonuniform level separations in this model, the actual SNR for transitions between the the closest two levels in this case is only 2.67. The relatively poor performance of smBEVO for this condition suggests that an SNR of approximately three or more is needed to reliably resolve neighboring levels. Preprocessing to denoise these time series may improve subsequent performance of smBEVO as high frequency information is generally not necessary to reliably resolve slower baseline drift. Although baseline estimation in all other tested conditions was overall excellent, there were a few outliers where the estimated and simulated baselines deviated significantly (see extreme MSE outliers in Fig. 2b and Supplemental Figs. 3, 5, 7 and 9). These extreme outliers represent time series for which the selected parameters for all 100 series were suboptimal. We note that in these cases two adjacent levels were often erroneously stitched together as the levels are built from the image trajectory segments, an aspect of the algorithm that could be improved upon in the future. One option is to simply remove these series from the dataset as they are readily identified by visual inspection. Alternatively, one could manually adjust the parameters for each of these cases individually to obtain a more visually compelling baseline estimate. Either approach should be feasible given their relative rarity. Of course, it is also possible to use a larger test set of series to initially identify optimal parameters, which may further reduce the already infrequent occurrence of poor estimations. Most importantly, for each unique combination of model, SNR, and drift type, the ability of smBEVO to produce accurate baseline estimations for 98-100 out of 100 time series based on minimal supervised selection of a single set of intuitive parameters (Supplementary Table 1) illustrates its practical relevance to baseline drift removal in large single-molecule datasets.

**Fig 3.**
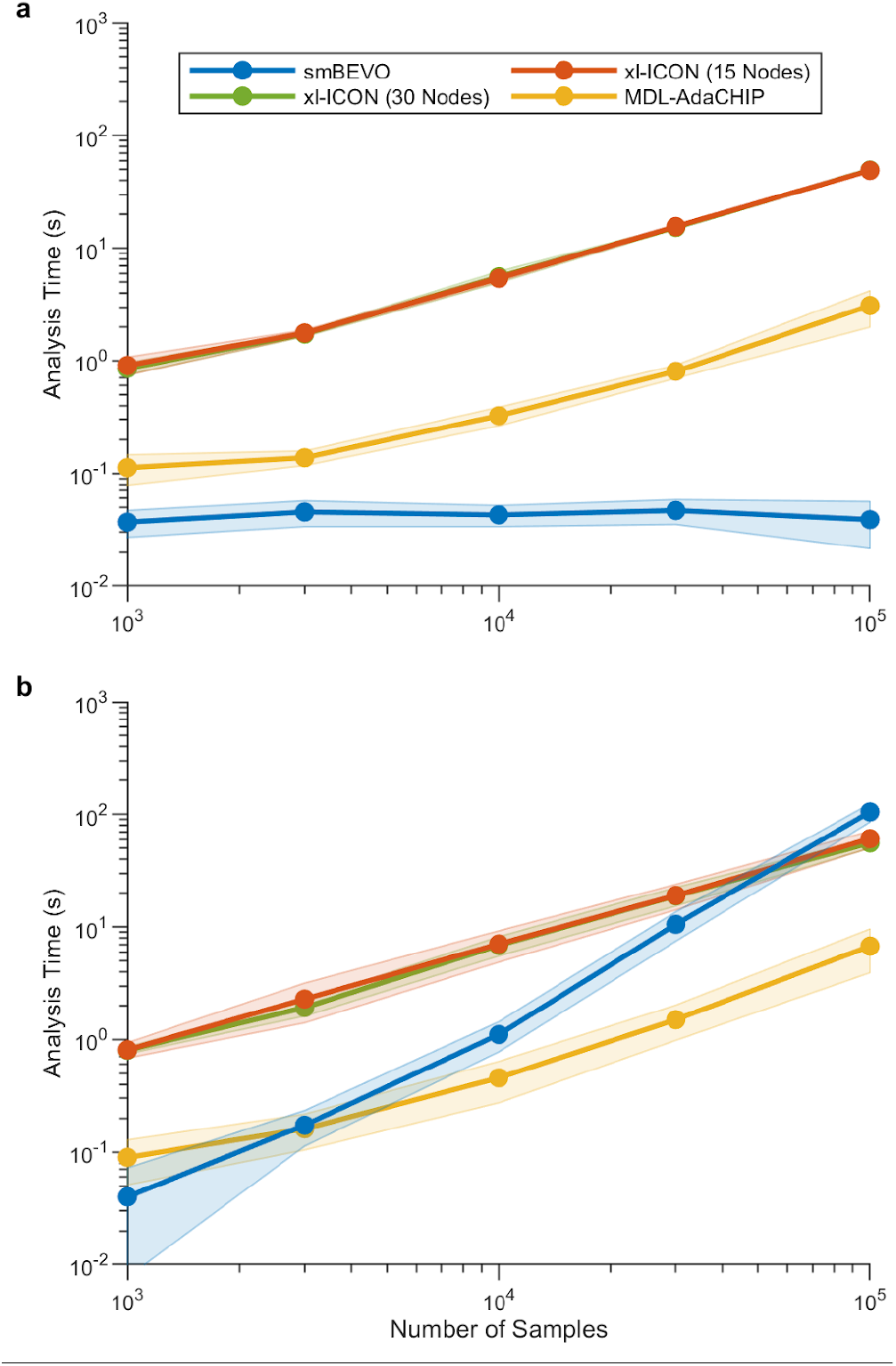
Computational speed of smBEVO depends on baseline rate of change rather than series length or sample frequency. **a-b**, Computation time as a function of series length for simulated data from the three-state cyclic model shown in Fig. 2 with added linear plus sinusoidal drift at SNR=6 (see Methods). Line and shaded region are mean ± standard deviation for ten series at each length. The sinusoidal drift period was either scaled such that three cycles spanned the series regardless of the number of samples (a) or held constant at three cycles per 1000 samples (b). Note that computation time for smBEVO depends primarily on the image resolution (i.e. *σ*_*x*_ and *σ*_*y*_) which itself depends on the drift frequency rather than the number of samples or sample frequency. For three cycles per series, smBEVO can use roughly the same sized pixel representations of each series regardless of the number of samples.

### Comparison of smBEVO with other baseline estimation methods

To better place smBEVO in context we compare its baseline estimation for the entire dataset described above to two other approaches for single-molecule drift correction: xl-ICON and MDL-AdaCHIP (Ouqamra & Bouilly, 2019; Sgouralis et al., 2018). Both approaches are unsupervised and do not require postulation of a specific model. xl-ICON utilizes iHMMs coupled with a node-based description of a time-varying baseline that is evolved to identify both drift and signal. In contrast, MDL-AdaCHIP is a signal compression approach combining the Minimum Description Length (MDL) principle with adaptive piecewise cubic Hermite interpolation to iteratively decompose the signal from confounding drift. We chose not to evaluate manual placement of baseline nodes as this approach is tedious and potentially infeasible for large datasets. Furthermore, estimated baselines via smBEVO were typically as good as anything we would have manually applied. We also chose not to evaluate optimization of a specific HMM coupled with baseline nodes (Venkataramanan & Sigworth, 2002). Although this approach may provide good results where applicable, postulation of a specific model can be nontrivial, and its application to high-throughput datasets where the behavior of interest occurs only in a subset of molecules may result in artifactual results for locations exhibiting alternative behaviors. Also, for camera-based imaging data, spatial nonuniformities in the optical pathways can result in per molecule variation in time series signal-to-noise that challenges global analysis by an HMM with specific state emissions. Thus, we limited our comparison to unsupervised approaches amenable to model-agnostic analysis. Although smBEVO is not completely unsupervised, user selection of parameters for this analysis was limited to visual inspection of a handful of series, and thereafter applied to all 100 series under the same conditions in an unsupervised manner. Furthermore, smBEVO is independent of any specific model and its parameters can be largely intuitively estimated, and thus its comparison to xl-ICON and MDL-AdaCHIP is relevant.

We applied xl-ICON and MDL-AdaCHIP to the same dataset as smBEVO using default parameters for both methods. Of the three methods, smBEVO provides the most visually compelling and accurate baseline estimates (i.e. lowest MSE between simulated and estimated baselines) across nearly all tested models, SNRs, and types of drift (Fig. 2, Supplementary Figs. 2-9). The only exception is for the three-state cyclic model with sinusoidal drift at an average SNR=4 as discussed above. Based on MSE scores xl-ICON was consistently the closest competitor. For control conditions without any applied drift xl-ICON performs similarly to smBEVO. However, xl-ICON’s performance is noticeably worse in the presence of baseline drift, at least in part due to its tendency to account for changes in baseline by adding additional artifactual states to its iHMM rather than adjusting the baseline nodes. This is exacerbated when the drift contains a linear component, making xl-ICON especially unsuited for data with linear drift. Adding additional baseline nodes (15 or 30 nodes per series) did not change this behavior. In contrast to xl-ICON, MDL-AdaCHIP has the opposite tendency to treat signal transitions between states as baseline drift. This tendency is exacerbated for series with long excursions in levels above the baseline, and is particularly evident for monotonic stepwise series where it may decide to treat the entire trend as baseline rather than a stepwise signal (e.g. Fig. 2a bottom). This makes MDL-AdaCHIP inappropriate for data such as that obtained from stepwise bleaching of fluorophores or largely monodirectional movement of molecular motors or enzymes along one-dimensional filaments or DNA strands (Heller et al., 2014; Myler et al., 2017; H. Zhang & Guo, 2014). Notably, this overfitting of the baseline variation can occur even in the absence of any added drift, albeit to a lesser extent.

In summary, smBEVO provides clearly superior overall baseline estimation as compared to xl-ICON and MDL-AdaCHIP across nearly all tested models, SNRs and types of drift. Strikingly, MSE between simulated and estimated baselines was nearly uniformly minimal independent of the type of drift applied (Fig. 2b; Supplemental Figs. 1-8). We acknowledge, however, that whereas xl-ICON and MDL-AdaCHIP are completely unsupervised approaches, smBEVO requires supervised selection of reasonable parameters based on a test set. Nonetheless, parameters are mostly intuitively estimated and for each unique combination of model, drift type and SNR we readily identified parameters that provided visually compelling baseline correction for nearly all simulated time series in the dataset.

### Computational speed

The computational cost of smBEVO depends primarily on the size of the data’s image representation rather than the length of the series. To resolve slow baseline variation in a small set of discrete levels, images on the order of a few hundred rows by a few hundred or a few thousand columns are often all that is required regardless of sample frequency. For such data, smBEVO is extremely efficient, taking only a few hundred milliseconds per series on a standard laptop with an Intel 7th generation core i7 processor (Fig. 3a). Not only is this much faster than xl-ICON and MDL-AdaCHIP, but it does not scale with sample frequency (Fig. 3a). However, for drift at a given average frequency, increasing series length at the same sample frequency will require larger images to resolve the extended baseline trend, which will increase computational cost (Fig. 3b). For example, relatively rapid baseline fluctuations on the millisecond time scale in single channel recordings lasting tens of minutes would require extremely large and costly image representations. Alternatively, if the drift is slow, then a much smaller image (i.e. relatively large *σ*_*x*_) will be needed even for high sample frequencies of 20 kHz or more. Keeping this limitation in mind, for many single-molecule datasets smBEVO will provide very efficient baseline estimation.

### Software

smBEVO is available as a MATLAB application that can be installed via the MATLAB interface. Source code and user manual are also available at: https://github.com/marcel-goldschen-ohm/smBEVO.

## Discussion

We present an efficient and accurate image-based approach to baseline correction for single-molecule time series, smBEVO. The approach is model-free and utilizes tools from image processing and computer vision to estimate the time-varying baseline drift that can in practice be difficult to eliminate in single-molecule experimental modalities. Given that nearly all analysis methods such as changepoint identification and HMMs rely heavily on the amplitude of the measured signal, removal of any baseline drift not associated with the behavior of the molecule of interest is a necessity for any analysis pipeline.

We show that smBEVO provides visually and quantitatively compelling baseline estimation for simulated data with multiple types of mild to aggressive drift (Fig. 2). We further applied smBEVO to several types of experimental single-molecule time series with noticeable drift such as the stepwise bleaching of fluorescence from GFP molecules or current during gating between closed and open conductance states of a single ion channel (Fig. 4). Although we do not have a ground truth to compare to as we did for the simulated data, it was generally very easy to rapidly obtain visually compelling baseline corrections that allow subsequent amplitude-based modeling of these data. In comparison to two other unsupervised approaches for drift correction, smBEVO was nearly always the most accurate while delivering the fastest computational speed (Figs. 2-3; Supplementary Figs. 2-9). Although not completely unsupervised, the approach relies on only a few key parameters that are largely intuitively estimated. Furthermore, given the fast execution time a reasonable parameter space around initial estimates can be efficiently explored. Finally, the same parameters are likely to be appropriate for nearly all series in a dataset collected under the same experimental conditions. Thus, smBEVO enables efficient, accurate and intuitive model-free baseline correction for many single-molecule datasets.

**Fig 4.**
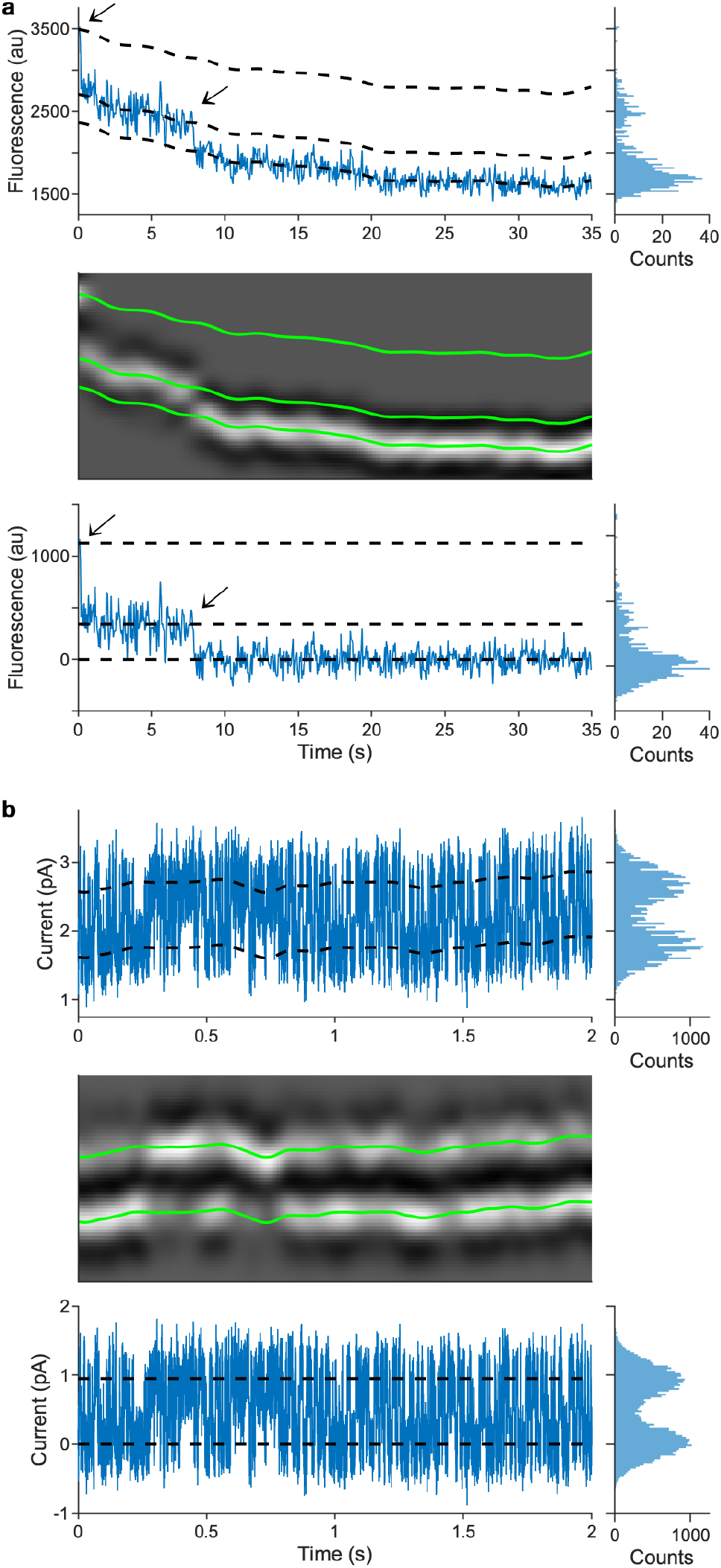
smBEVO applied to experimental data. **a**, top) TIRF imaging of the stepwise bleaching of two GFP molecules (arrows) contaminated by a slow background decay. Dashed lines are levels identified by smBEVO. Parameters are *σ*_*x*_= 0.7 sec, *σ*_*y*_= 150 au, *λ*= 2, *α*= 1, and *β*= 1. middle) Image representation of the data overlaid with identified levels (green lines). bottom) Baseline subtracted data series. **b**, top) Recording of current through a single ion channel with predominant gating between two different conductance states. The baseline current is contaminated by a slow wobble possibly due to the stability of the lipid membrane patch in the recording pipet. Dashed lines are levels identified by smBEVO. Parameters are *σ*_*x*_= 0.035 sec, *σ*_*y*_= 0.2 pA, *λ*= 0.5, *α*= 1, and *β*= 1. middle) Image representation of the data overlaid with identified levels (green lines). bottom) Baseline subtracted data series.

## Online Methods

### Image representation of the data series

Results are highly dependent on *σ*_*x*_ and *σ*_*y*_, which together with *s*_*x*_ and *s*_*y*_ define the image resolution of the data series. However, increasing *s*_*x*_ and *s*_*y*_ above 4-8 pixels has relatively little impact on baseline estimation, and for optimal computational speed should be kept as small as possible such that the series behavior is still resolved in the image. In practice, *s*_*x*_= *s*_*y*_= 4 pixels typically works well. The image representation of the data series is obtained as a normalized two-dimensional histogram of the series (*x, y*) data points with bin widths in each dimension of *σ*_*x*_/*s*_*x*_ and *σ*_*y*_/*s*_*y*_. Actual bin widths along the x-axis may be slightly reduced to ensure an integer number of bins span the series. Bin edges along the y-axis are separated by *σ*_*y*_/*s*_*y*_ and centered with respect to the data y-range. Prior to normalization, the value in each bin is the number of data samples that fall within the bin. To avoid edge artifacts during subsequent filtering, the top and bottom of the image is padded with *ceil*(5*s*_*y*_) rows of zeros.

### Image filter

The image filter kernel *k*_*x,y*_ is proportional to a Gaussian along the x-axis and the second derivative of a Gaussian along the y-axis with standard deviations of *s*_*x*_ and *s*_*y*_ pixels, respectively (Eqs. 1-2, Fig. 1c inset).

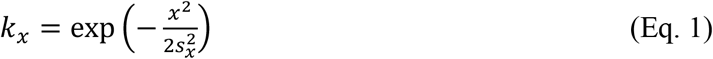

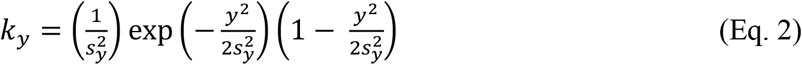

This filter was previously used to detect lane markers in computer vision-based approaches to automated driving (Aly, 2008). Here we repurpose it to enable detection of distinct levels in single-molecule time series.

### Initial image trajectory segmentation

The maximum intensity pixel in each column provides an initial denoised trajectory in image coordinates that approximates the data series trend while largely maintaining sharp jumps between amplitude levels (Fig. 1d). However, there can occur instances where steep jumps are not maintained. In these cases, a distinct jump is often identified in the analogous conjugate trajectory for the minimum intensity pixel in each column (Fig. 1d). Thus, potential changepoints are identified as locations where jumps in either the maximum or minimum trajectories exceed a specified threshold of *s*_*y*_/2. The maximum intensity trajectory is divided into disjoint segments at the identified changepoints. Furthermore, to limit skew at the ends of each segment that can occur due to smoothing by the filter kernel, the pixels corresponding to the first and last *ceil*(*s*_*x*_/2) columns of each segment are replaced with pixels at the same row index (corresponding to amplitude in the series) as the immediately adjacent interior pixels in the segment. Segments shorter than 2*ceil*(*s*_*x*_/2) columns are left as is.

### Simulated single-molecule time series

All simulations and analyses were conducted in MATLAB version R2021a (Mathworks). Single-molecule time series were simulated as Markov chains consisting of dwells in distinct states. The four simulated Markov mechanisms are depicted in Fig. 2a where each state is labeled with its mean emission amplitude and allowed transitions between states are indicated by unidirectional or bidirectional arrows. All simulations had a uniform sample frequency *f*_*s*_. For the two-state model both forward and backward transition rates were set to 0.01*f*_*s*_. For the three-state cyclic model transition rates between the two highest amplitude states were set to 0.01*f*_*s*_ and all other rates were set to 0.003*f*_*s*_. For the four- and five-state linear models all transition rates were set to 0.005*f*_*s*_. These prescriptions provide simulations that test performance on both equivalent and disparate rates within a given model. To simulate noisy experimental data, Gaussian noise was added to the noiseless series described above. The standard deviation of the added noise (*σ*) was set according to a desired average signal-to-noise ratio such that *SNR*= Δ*I*_*avg*_/*σ*, where Δ*I*_*avg*_ is the average separation between mean emission amplitudes of neighboring states. Five types of baseline drift were added to the noisy time series to simulate various empirically observed drifts: no drift, linear, sinusoidal, linear plus sinusoidal, and exponential. Linear drift was applied with a slope of 0.1 au per 1000 samples. Sinusoidal drift had an amplitude of 0.15 au and a frequency of three cycles per 5000 samples. These two drifts were summed together to generate linear plus sinusoidal drift. Exponential drift was applied with an amplitude of 0.5 au and a time constant of 1000 samples.

## Supplementary Material

**Supplementary Fig 1.**
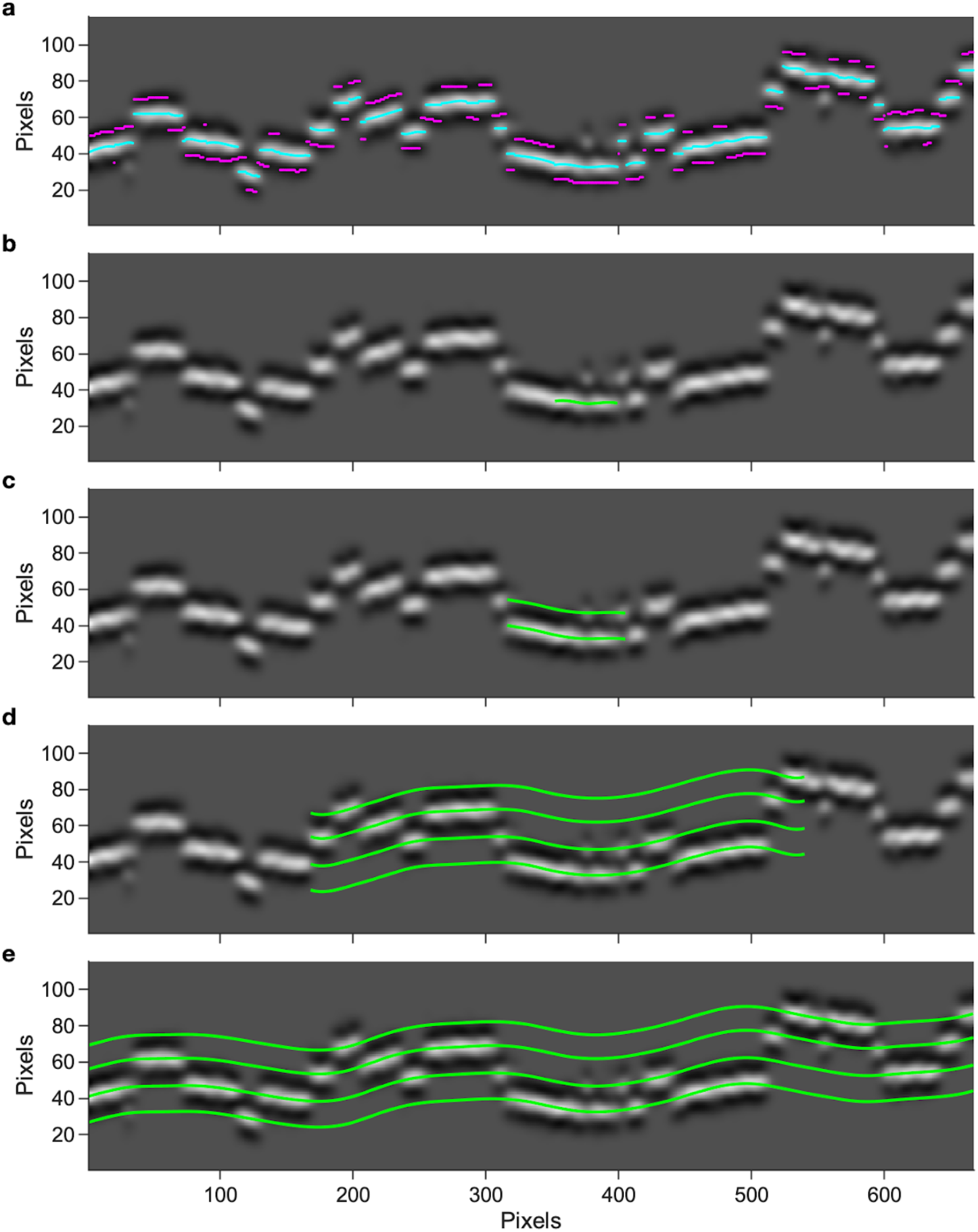
Illustration of method for building levels from the denoised image trajectory. **a**, Image of time series simulated with the four-state linear model shown in Figs. 2 and S6 with added linear and sinusoidal drift at SNR=4 after filtering with *σ*_*x*_ = 30 and *σ*_*y*_ = 0.1. Trajectories for the maximum (cyan) and minimum (magenta) intensity pixels in each column are shown broken into segments as described in the main text. **b**, The longest maximum intensity segment between jumps in either the maximum or minimum intensity trajectories (green) is used to initialize the first amplitude level in the image. **c-e**, Snapshots of the process of iteratively extending the current set of levels into the adjacent segments on either side. If the adjacent maximum intensity segment is sufficiently different from all the current levels, a new level is added and back extrapolated along the current levels. This process is repeated until the identified levels span the entire image. Levels were smoothed at each iteration according to *λ* = 7 and a minimum level separation of *δ*_*y*_ = 0.1 was enforced.

**Supplementary Fig. 2.**
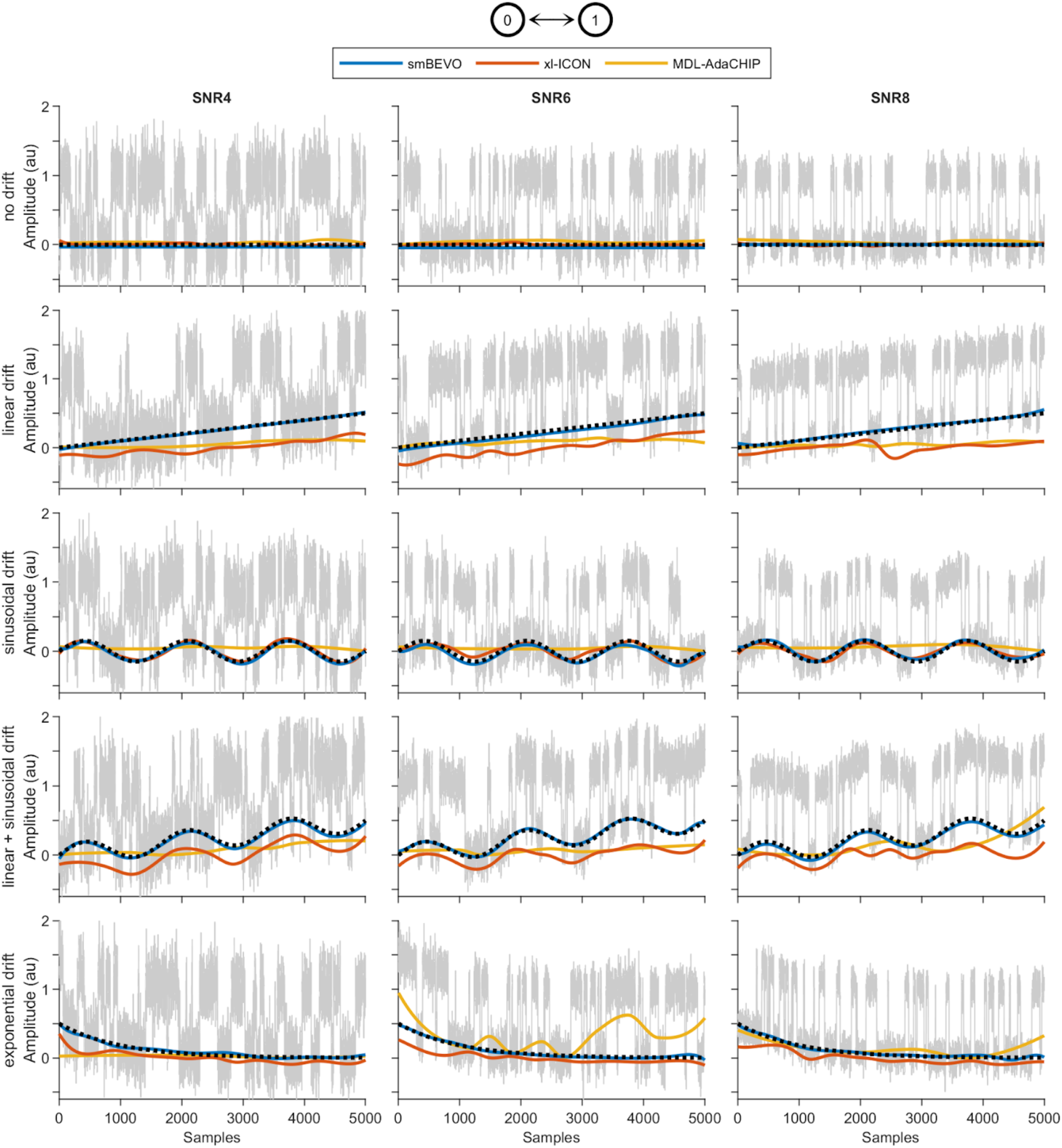
Examples of simulated data for a two-state model with drift. Simulated noisy time series for a two-state model with state emission amplitudes of 0 and 1 (top, see Methods). Each row has a different type of added baseline drift (dotted) and each column a different simulated signal-to-noise ratio (SNR). Series are overlaid with baseline estimations using smBEVO, xl-ICON or MDL-AdaCHIP.

**Supplementary Fig. 3.**
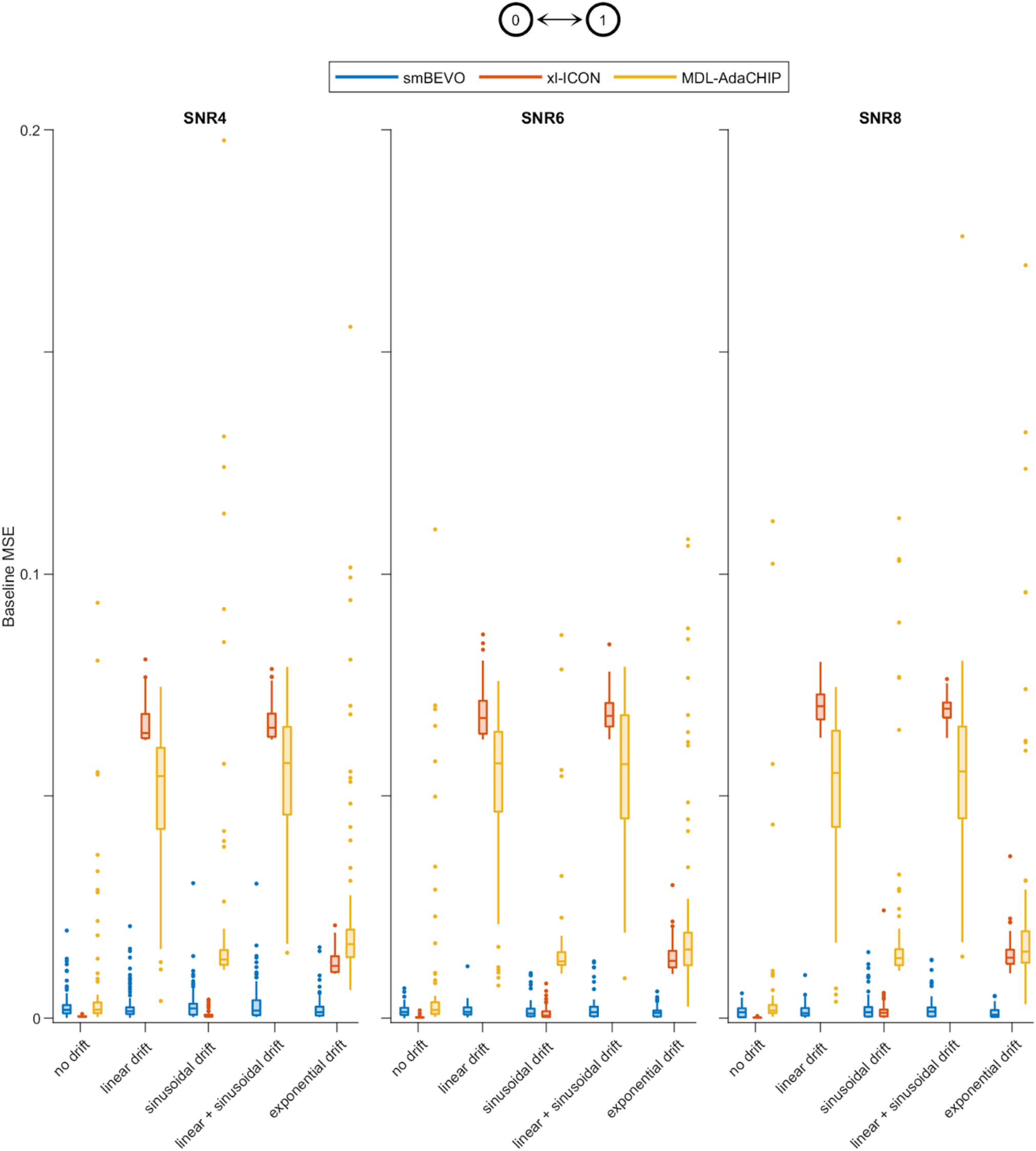
Summary of baseline estimation for a two-state model with drift. Box plots for the mean squared error (MSE) between simulated and estimated baselines for 100 simulated series as shown in Supplementary Fig. 1 at each unique combination of SNR and drift type. Dots are outliers. For MDL-AdaCHIP two outliers with MSEs greater than 0.2 are not shown.

**Supplementary Fig. 4.**
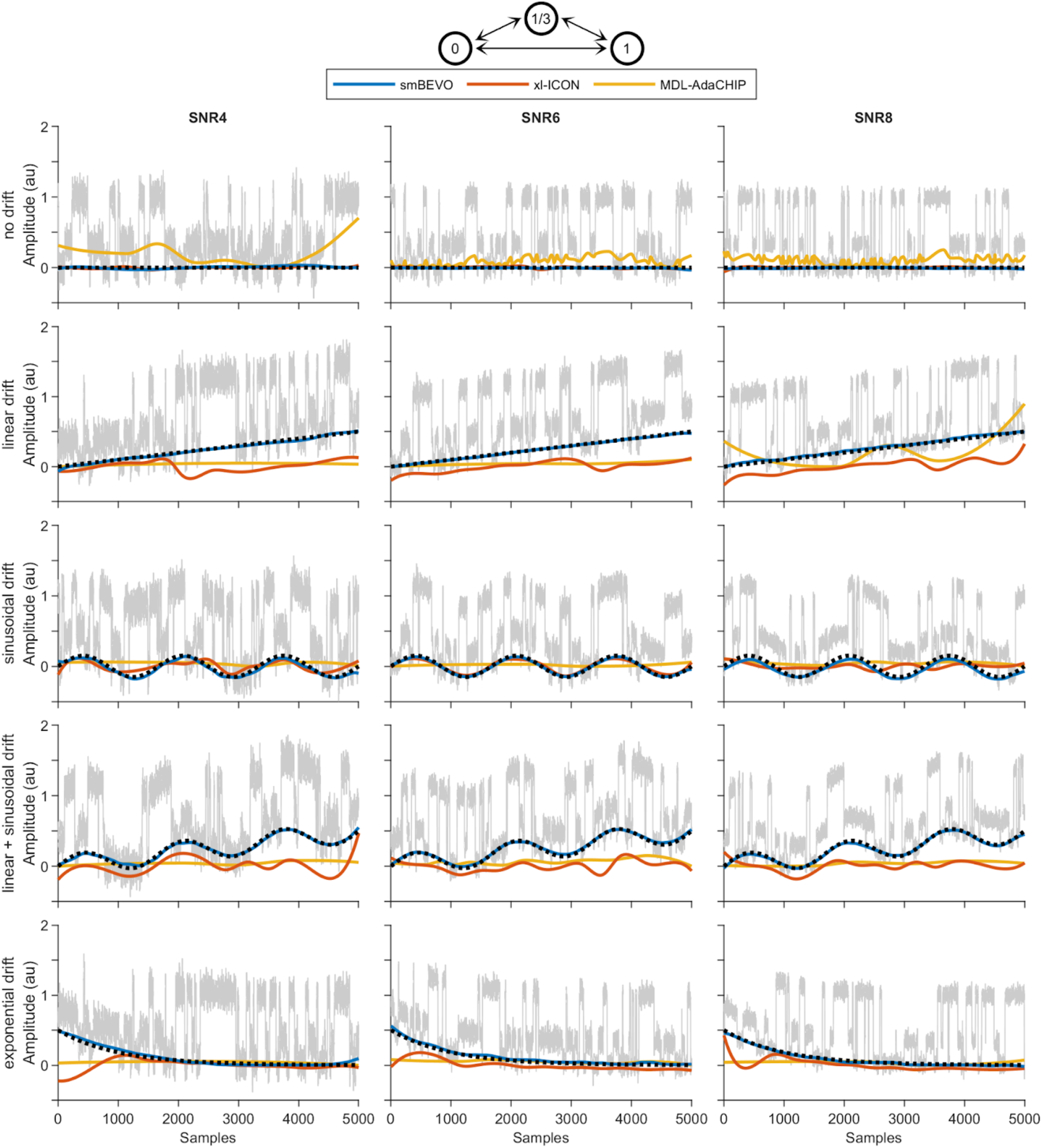
Examples of simulated data for a three-state cyclic model with drift. Simulated noisy time series for a three-state cyclic model with state emission amplitudes of 0, ⅓ and 1 (top, see Methods). Each row has a different type of added baseline drift (dotted) and each column a different simulated signal-to-noise ratio (SNR). Series are overlaid with baseline estimations using smBEVO, xl-ICON or MDL-AdaCHIP.

**Supplementary Fig. 5.**
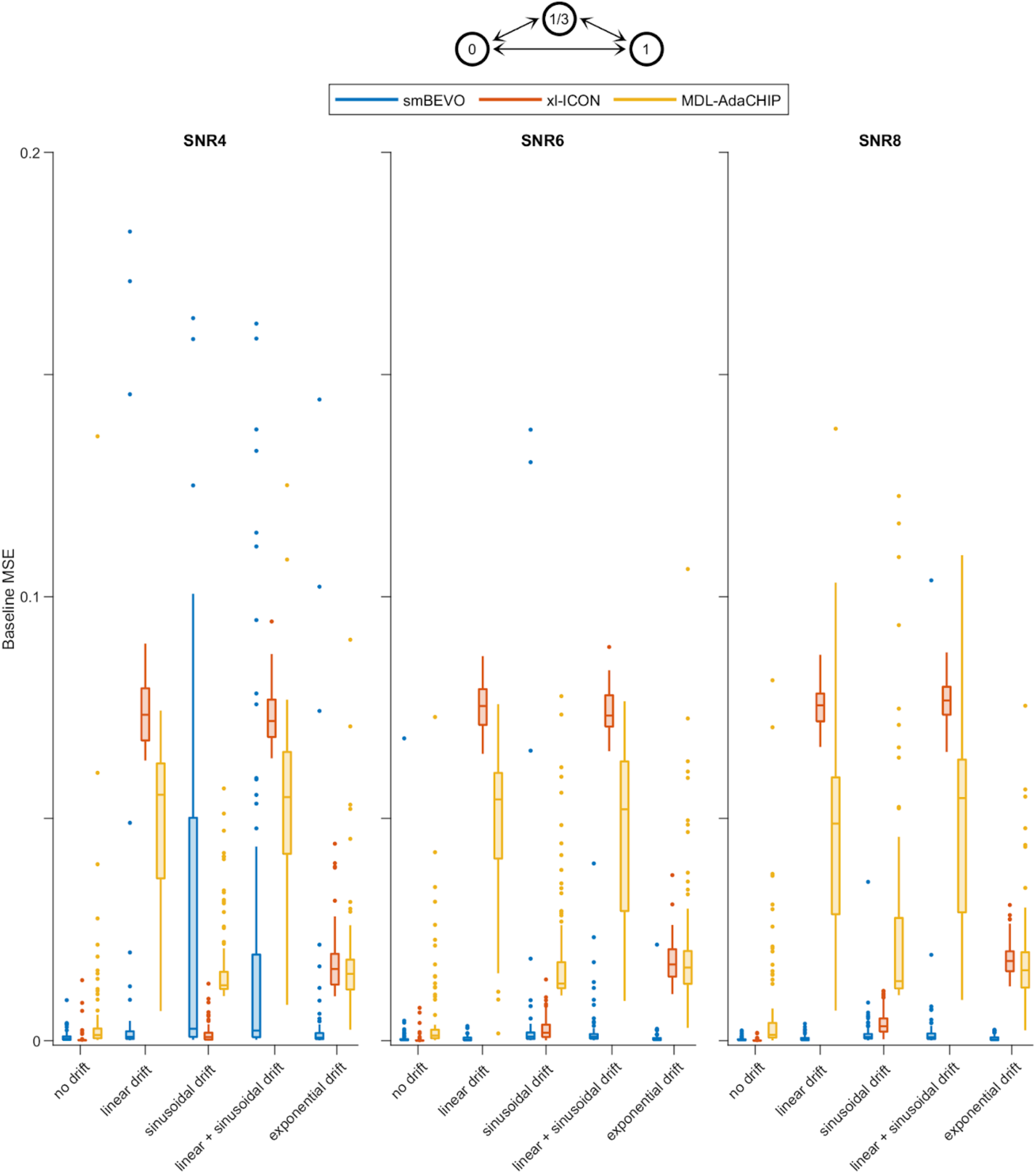
Summary of baseline estimation for a three-state cyclic model with drift. Box plots for the mean squared error (MSE) between simulated and estimated baselines for 100 simulated series as shown in Supplementary Fig. 3 at each unique combination of SNR and drift type. Dots are outliers. For smBEVO and MDL-AdaCHIP 13 or two outliers with MSEs greater than 0.2 are not shown, respectively.

**Supplementary Fig. 6.**
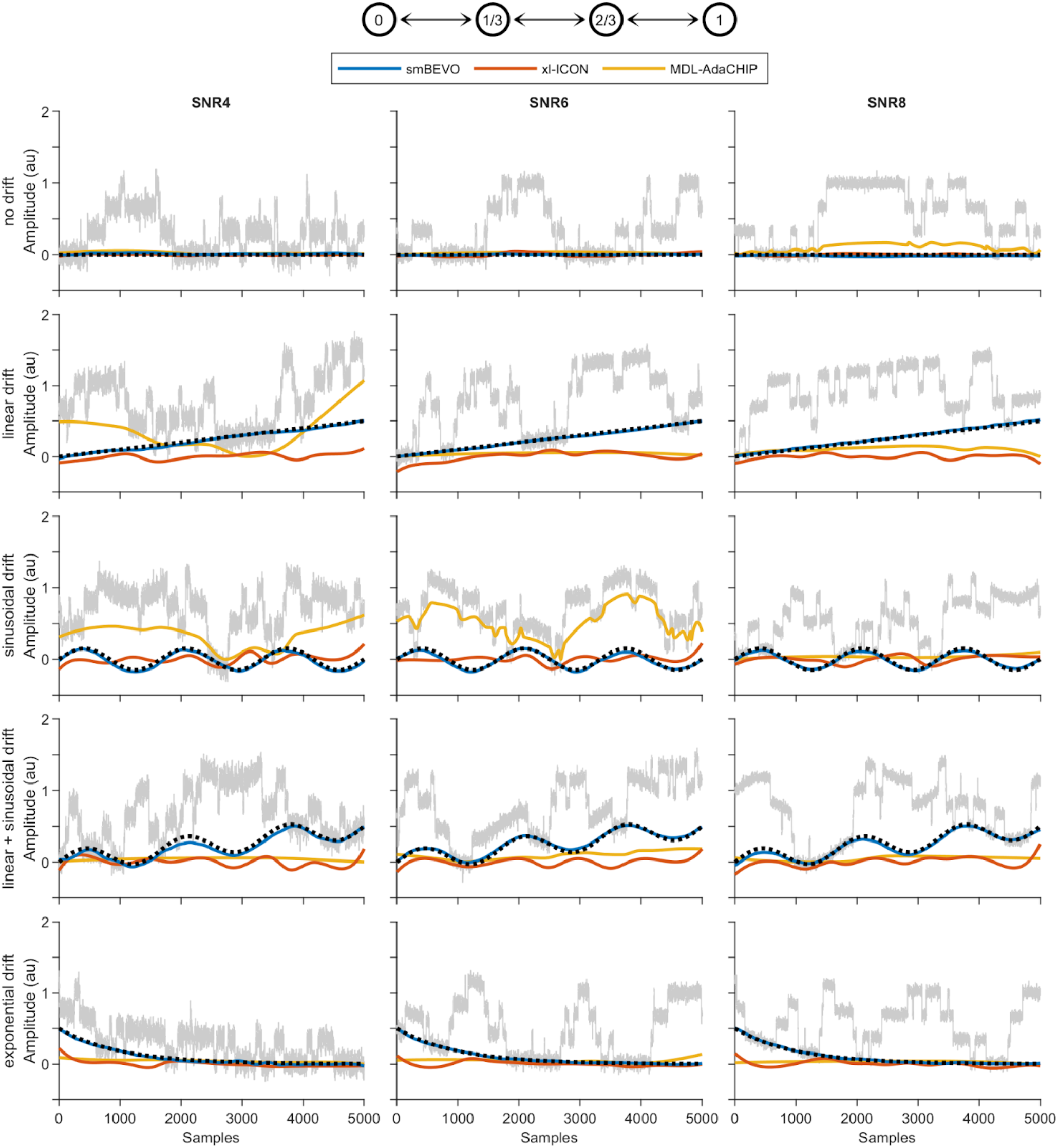
Examples of simulated data for a four-state linear model with drift. Simulated noisy time series for a four-state linear model with state emission amplitudes of 0, ⅓, ⅔ and 1 (top, see Methods). Each row has a different type of added baseline drift (dotted) and each column a different simulated signal-to-noise ratio (SNR). Series are overlaid with baseline estimations using smBEVO, xl-ICON or MDL-AdaCHIP.

**Supplementary Fig. 7.**
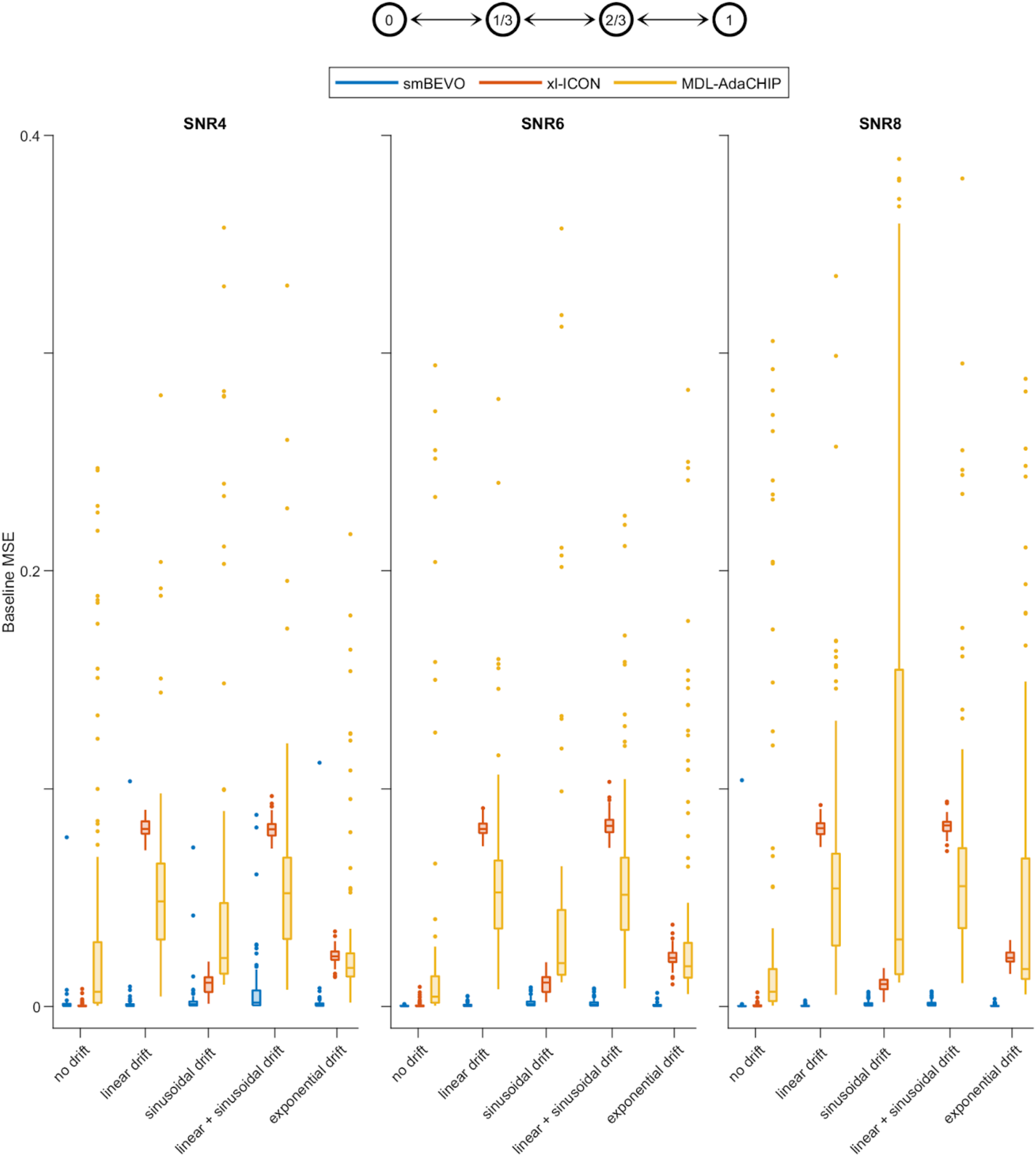
Summary of baseline estimation for a four-state linear model with drift. Box plots for the mean squared error (MSE) between simulated and estimated baselines for 100 simulated series as shown in Supplementary Fig. 5 at each unique combination of SNR and drift type. Dots are outliers. For MDL-AdaCHIP five outliers with MSEs greater than 0.4 are not shown.

**Supplementary Fig. 8.**
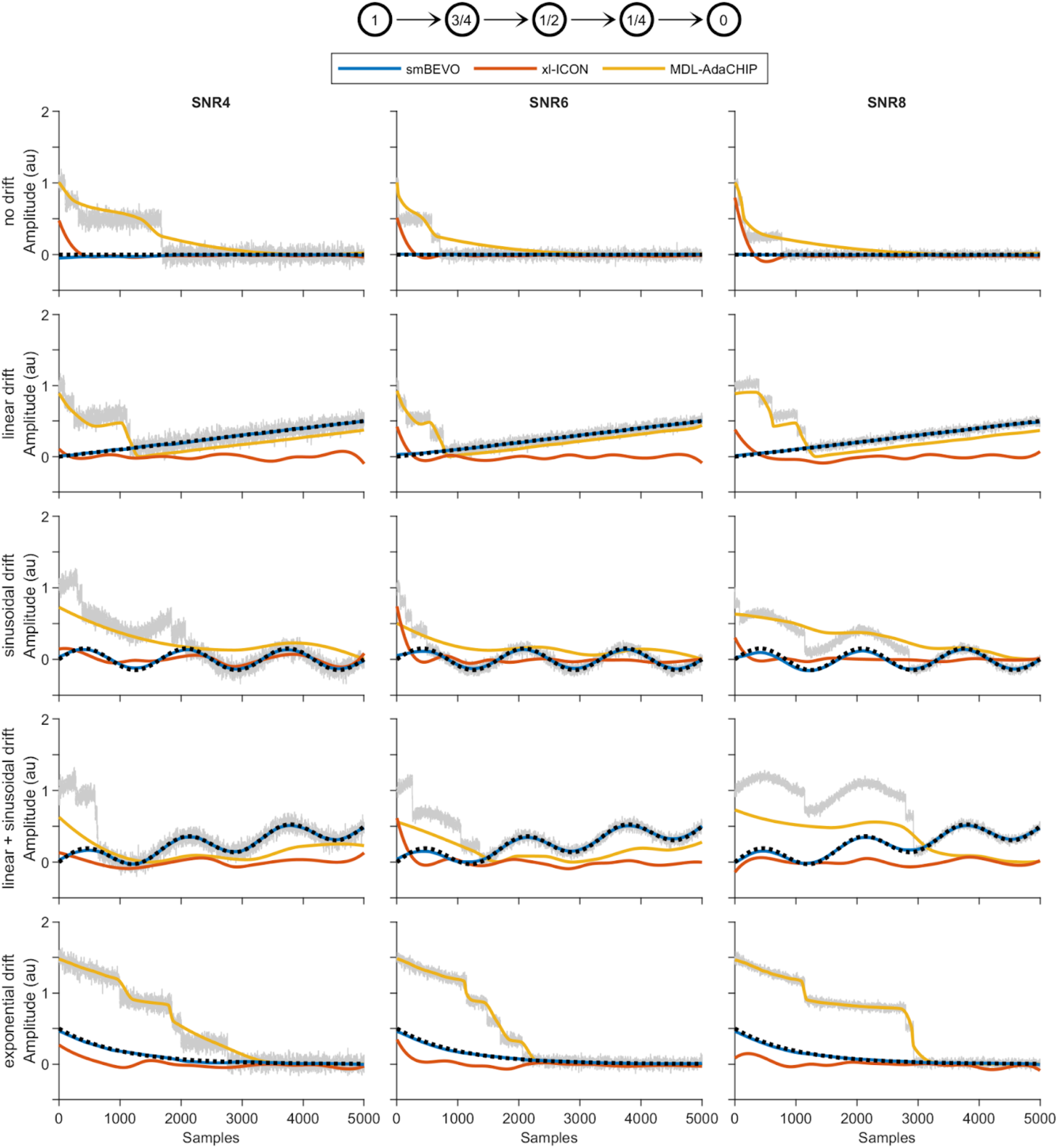
Examples of simulated data for a five-state unidirectional linear model with drift. Simulated noisy time series for a five-state unidirectional linear model with state emission amplitudes of 1, ¾, ½, ¼ and 0 (top, see Methods). Each row has a different type of added baseline drift (dotted) and each column a different simulated signal-to-noise ratio (SNR). Series are overlaid with baseline estimations using smBEVO, xl-ICON or MDL-AdaCHIP.

**Supplementary Fig. 9.**
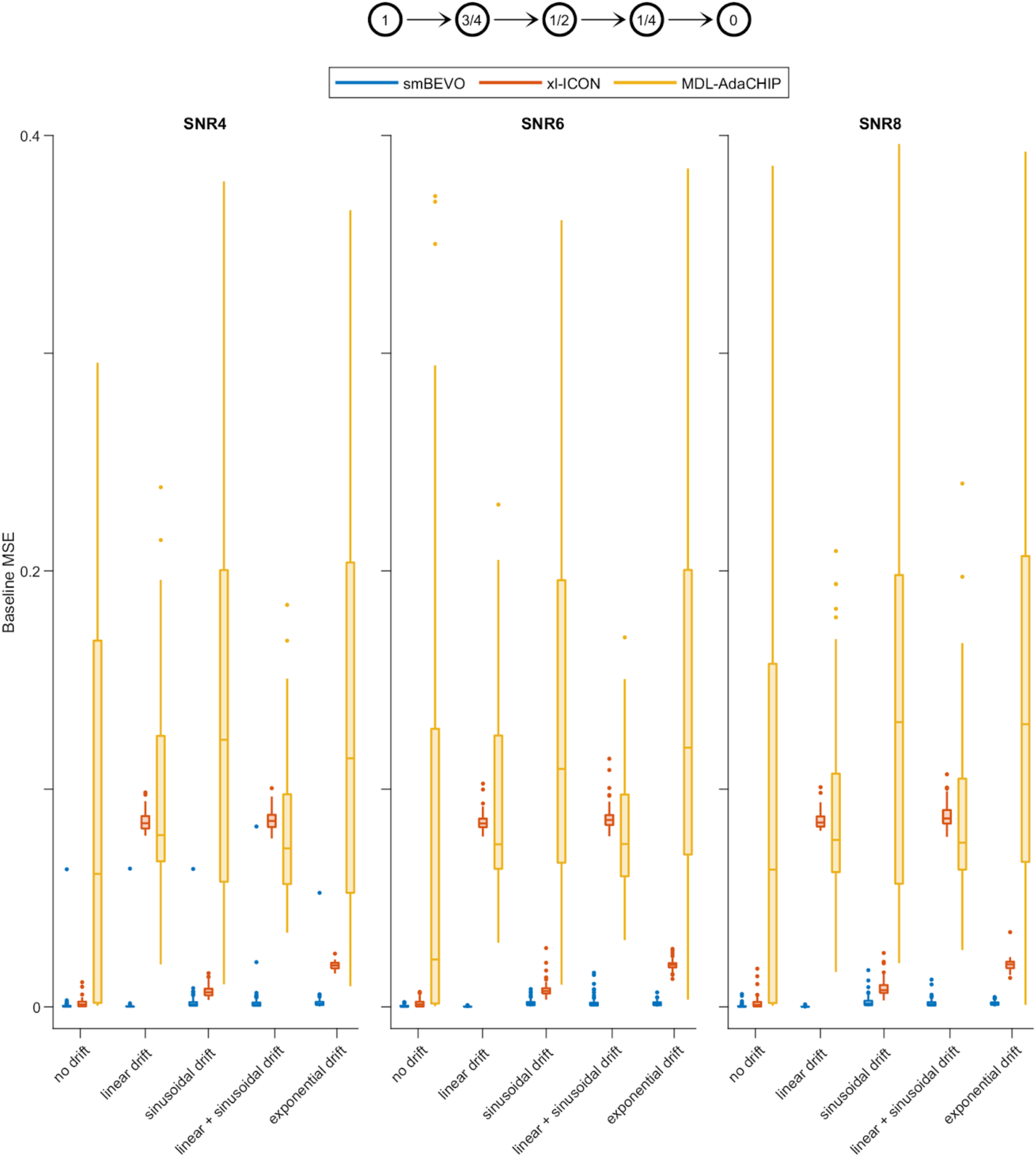
Summary of baseline estimation for a five-state unidirectional linear model with drift. Box plots for the mean squared error (MSE) between simulated and estimated baselines for 100 simulated series as shown in Supplementary Fig. 7 at each unique combination of SNR and drift type. Dots are outliers. For MDL-AdaCHIP 13 outliers with MSEs greater than 0.4 are not shown.

**Supplementary Table 1.** smBEVO parameters used for simulations of the two-state model with various types of baseline drift. For this model, representative examples of smBEVO drift estimates are shown in Supplementary Figure 2, and summary of overall smBEVO results are shown in Supplementary Figure 3. Adjustment of the baseline estimate using active contours was not necessary at any SNR/Drift condition for this model. *Where not specified, a default value of 1.75*δ*_*y*_ was used.

|  | SNR = 4 |  |  |  | SNR = 6 |  |  |  | SNR = 8 |  |  |  |
| --- | --- | --- | --- | --- | --- | --- | --- | --- | --- | --- | --- | --- |
| Drift | $\sigma_x$ | $\sigma_y$ | $\lambda$ | $\delta_y^*$ | $\sigma_x$ | $\sigma_y$ | $\lambda$ | $\delta_y^*$ | $\sigma_x$ | $\sigma_y$ | $\lambda$ | $\delta_y^*$ |
| none | 300 | 0.33 | 6 | — | 350 | 0.33 | 6 | — | 200 | 0.33 | 6 | — |
| linear | 300 | 0.33 | 6 | — | 200 | 0.33 | 6 | — | 200 | 0.33 | 6 | — |
| sinusoidal | 60 | 0.33 | 6 | — | 60 | 0.33 | 6 | — | 60 | 0.33 | 6 | — |
| linear + sinusoidal | 60 | 0.33 | 6 | — | 60 | 0.33 | 6 | — | 60 | 0.33 | 6 | — |
| exponential | 60 | 0.33 | 8 | — | 60 | 0.33 | 8 | — | 60 | 0.33 | 8 | — |

**Supplementary Table 2.**
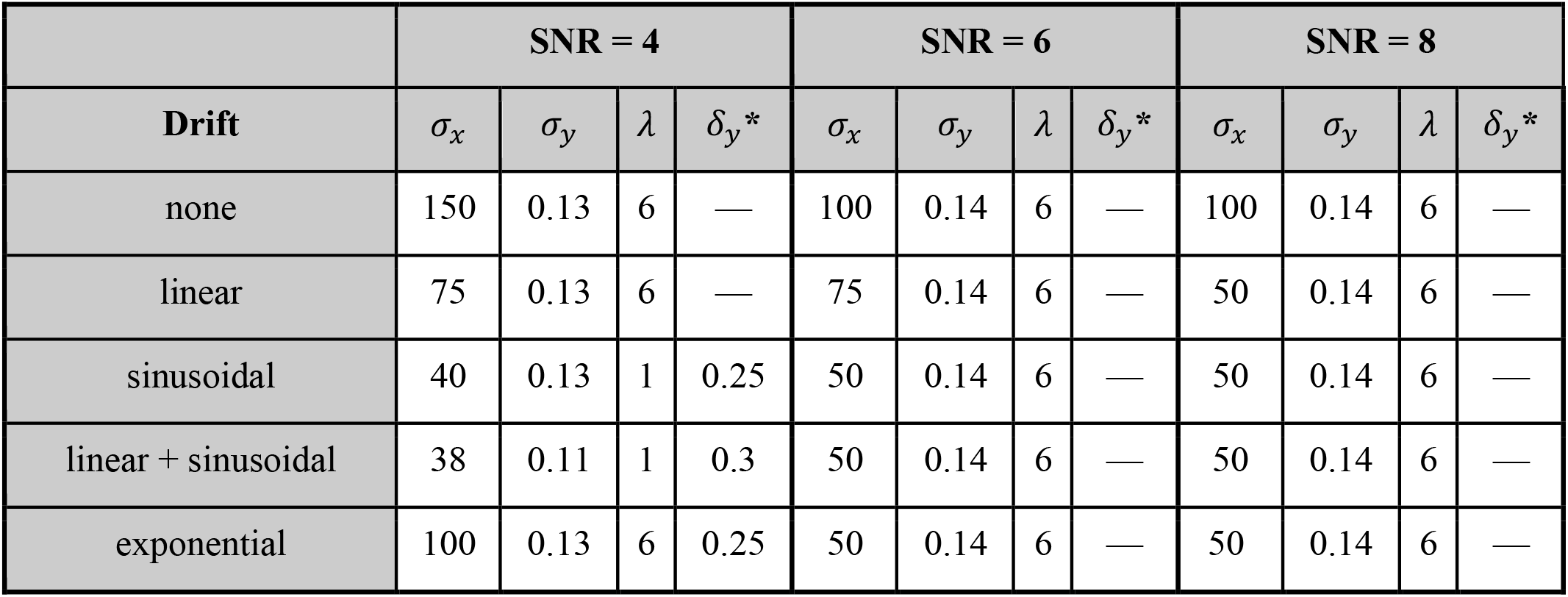
smBEVO parameters used for simulations of the three-state cyclic model with various types of baseline drift. For this model, representative examples of smBEVO drift estimates are shown in Supplementary Figure 4, and summary of overall smBEVO results are shown in Supplementary Figure 5. For SNR = 4, the linear and linear + sinusoidal cases required adjustment of the baseline estimate using active contours (sinusoidal: *α* = 5, *β* = 1; linear + sinusoidal: *α* = 7, *β* = 1). *Where not specified, a default value of 1.75*δ*_*y*_ was used.

**Supplementary Table 3.**
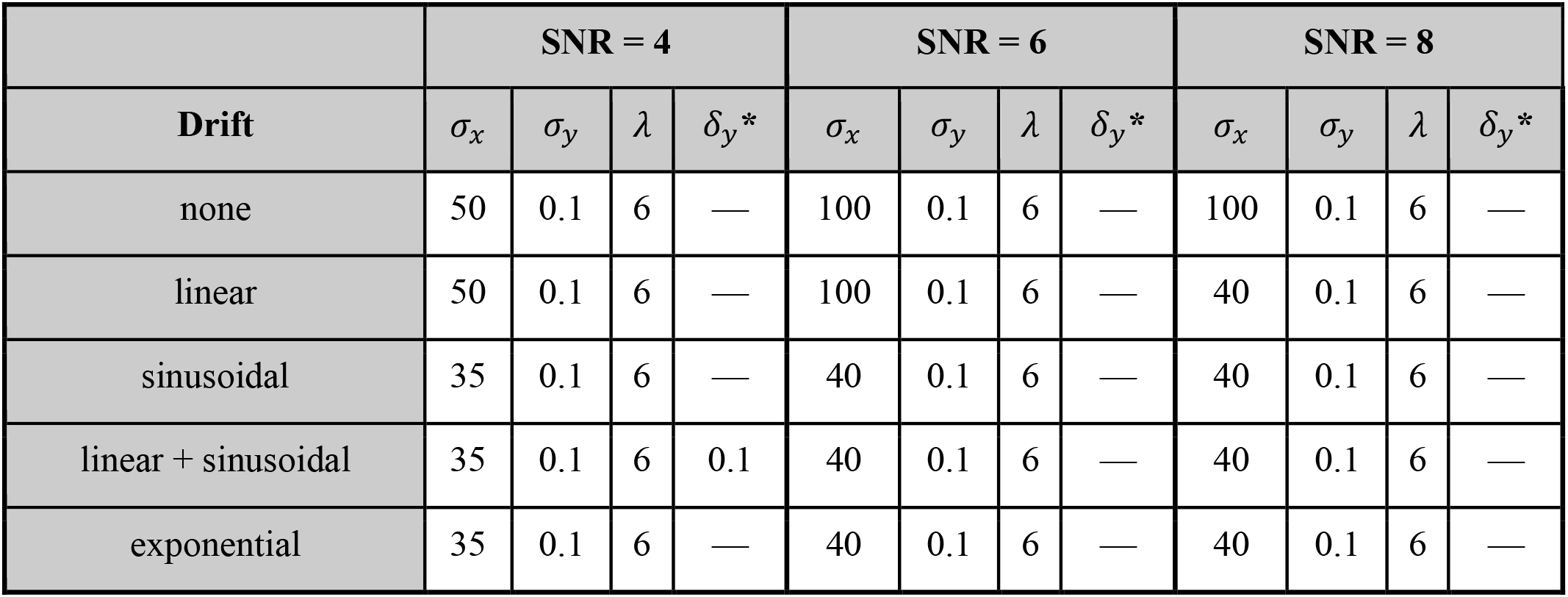
smBEVO parameters used for simulations of the four-state linear model with various types of baseline drift. For this model, representative examples of smBEVO drift estimates are shown in Supplementary Figure 6, and summary of overall smBEVO results are shown in Supplementary Figure 7. Adjustment of the baseline estimate using active contours was not necessary at any SNR/Drift condition for this model. *Where not specified, a default value of 1.75*δ*_*y*_ was used.

|  | SNR = 4 |  |  |  | SNR = 6 |  |  |  | SNR = 8 |  |  |  |
| --- | --- | --- | --- | --- | --- | --- | --- | --- | --- | --- | --- | --- |
| Drift | $\sigma_x$ | $\sigma_y$ | $\lambda$ | $\delta_y^*$ | $\sigma_x$ | $\sigma_y$ | $\lambda$ | $\delta_y^*$ | $\sigma_x$ | $\sigma_y$ | $\lambda$ | $\delta_y^*$ |
| none | 50 | 0.1 | 6 | — | 100 | 0.1 | 6 | — | 100 | 0.1 | 6 | — |
| linear | 50 | 0.1 | 6 | — | 100 | 0.1 | 6 | — | 40 | 0.1 | 6 | — |
| sinusoidal | 35 | 0.1 | 6 | — | 40 | 0.1 | 6 | — | 40 | 0.1 | 6 | — |
| linear + sinusoidal | 35 | 0.1 | 6 | 0.1 | 40 | 0.1 | 6 | — | 40 | 0.1 | 6 | — |
| exponential | 35 | 0.1 | 6 | — | 40 | 0.1 | 6 | — | 40 | 0.1 | 6 | — |

**Supplementary Table 4.** smBEVO parameters used for simulations of the five-state unidirectional linear model with various types of baseline drift. For this model, representative examples of smBEVO drift estimates are shown in Supplementary Figure 8, and summary of overall smBEVO results are shown in Supplementary Figure 9. Adjustment of the baseline estimate using active contours was not necessary at any SNR/Drift condition for this model. *Where not specified, a default value of 1.75*δ*_*y*_ was used.

|  | SNR = 4 |  |  |  | SNR = 6 |  |  |  | SNR = 8 |  |  |  |
| --- | --- | --- | --- | --- | --- | --- | --- | --- | --- | --- | --- | --- |
| Drift | $\sigma_x$ | $\sigma_y$ | $\lambda$ | $\delta_y^*$ | $\sigma_x$ | $\sigma_y$ | $\lambda$ | $\delta_y^*$ | $\sigma_x$ | $\sigma_y$ | $\lambda$ | $\delta_y^*$ |
| none | 100 | 0.1 | 3 | — | 150 | 0.1 | 3 | — | 150 | 0.1 | 3 | — |
| linear | 50 | 0.1 | 3 | — | 50 | 0.1 | 3 | — | 50 | 0.1 | 3 | — |
| sinusoidal | 40 | 0.1 | 3 | 0.15 | 50 | 0.1 | 3 | 0.15 | 50 | 0.1 | 3 | — |
| linear + sinusoidal | 40 | 0.1 | 3 | 0.15 | 50 | 0.1 | 3 | 0.15 | 50 | 0.1 | 3 | 0.15 |
| exponential | 50 | 0.1 | 3 | — | 50 | 0.1 | 3 | — | 50 | 0.1 | 3 | 0.15 |

